# A new genetic method to predict the prognostic outcome of melanoma

**DOI:** 10.1101/2020.03.09.984732

**Authors:** Hong GuoHu, Guan Qing, Luo XinHua

## Abstract

Cutaneous melanoma is quite often encountered in dermato-oncology. This paper describes a new genetic method to predict the prognostic outcome of melanoma. Data were collected from the TCGA databases. According to tumor progression status, the data were divided into two groups to evaluate the significant biological processes and key genes influencing the outcome of melanoma using a bioinformatics method. By adopting a statistical regression analysis method, a novel score based on the contributing genes was developed. Cox regression analysis was used to validate the effectiveness of the genetic risk score in predicting the outcome. Seven biological processes associated with melanocytes were identified. A protein-protein interactions network showed that 27 functional genes were associated with the outcome of melanoma. Among these, three genes (COL17A1, ITGA6, and SPRR2F) were used to calculate the genetic risk score, which was regarded as an independent and effective risk factor for disease progression or overall survival in melanoma.

## Introduction

The incidence of melanoma has been increasing rapidly in recent years and has become an important health problem globally^(1, 2)^. Melanoma can be treated with satisfactory results by surgical resection if detected early. However, with tumor metastasis or recurrence, the prognosis becomes very poor^(3)^. Early evaluation of the disease progression could effectively guide individualized treatment. The general prognostic assessment criteria are mostly based on clinical information^(3)^. Furthermore, genetic prognostic methods could be used to expand this particular research field, e.g., studies on BRAF^(4)^ mutation and its specific inhibitor have provided good results. Thus, screening of more specific genes as prognostic predictors in melanoma is of enormous importance.

In this study, the melanoma cohort data provided by the TCGA (The Cancer Genome Atlas)^(5)^ was used to identify the significant biological processes and key genes for predicting the outcome of melanoma and thereby develop a new prognostic assessment method.

## Materials and methods

The TCGA database was used for obtaining data from a cohort of 469 patients with cutaneous melanoma. The normalized gene mRNA expression profile value and clinical data (age, sex, TNM stage, ulceration, and drug treatment) were downloaded from the cBioportal website (www.cbioportal.org)^(6)^.

In order to screen the functional genes affecting the tumor progression status, we defined rapid progression group and long-term stability group. The rapid progression group included 68 patients who had progressive or recurrent diseases within 1 year. The long-term stability group included 33 patients who had a disease-free status over 5 years and 72 patients who had progressive or recurrent diseases but were disease-free for least 5 years. We identified the differentially expressed genes (DEGs) in the rapid progression and long-term stability groups by calculating the log_2_ (FC) (fold-change) of mRNA expression value and adjusted the corresponding p-values by the FDR method with a threshold adjusted p-value of < 0.05 and |FC| value of > 0.5.

Enrichment analysis of DEGs was performed online with the Metascape tool (www.metascape.org). The useful biological functions were concentrated in the gene ontology biological process (GOBP) terms. According to the p-value and the relationship with the skin, we identified the topic GOBP terms. The genes enriching the topic terms were used to extract the functional genes. We constructed the protein-protein interaction (PPI) network of the enriched gene by Cytoscape software^(7)^ (3.7.1) and STRING app^(8)^. The combined score of > 0.4 was set as the cut-off criterion, and the genes included in this network were identified as the key functional genes associated with disease progression.

For identifying the markers to predict disease progression, we performed a statistical analysis of the appropriate patients. The data of 469 patients provided by the TCGA database were browsed, and the data of 372 patients were included (68 cases were excluded because of incomplete disease status information and 29 cases were excluded because of disease-free status with an observation period of < 1 year). The mRNA expression profile value was log-transformed (logarithm to the base 2) by the least absolute shrinkage and selection operator (LASSO) method^(9)^. The findings were cross-validated for three times to identify the method of selection of the candidate genes and constructing a prediction model for genetic risk score. We found an optimal cutoff genetic risk score by X-tile software^(10)^ in these 372 patients. We divided the target data into high-risk and low-risk groups based on the Kaplan-Meier survival analysis and log-rank test.

We evaluated the role of the genetic risk score in predicting the melanoma outcome both in terms of disease progression and overall survival. The data of 372 patients obtained previously were used to perform the analysis for disease progression. A total of 425 patients (9 cases with incomplete survival information and 35 living cases with an observation time of < 1 year were excluded) were used to analyze the overall survival. The clinical characteristics (age, sex, TNM stage, ulceration, and drug treatment) were downloaded. A univariate Cox regression analysis was used to calculate the hazard ratio (HR) and p-value among the clinical features and genetic risk score. We selected the statistically significant variables for performing multivariate Cox regression analysis to evaluate the independent and effective risk factors.

All statistical operations were performed with R software version 3.5.1. Limma package was used to identify the DEGs, and six mainstream packages (mnet, pec, rms, survminer, survIDINRI, and survival) were used to perform the statistical analyses. A p value of < 0.05 was considered statistically significant.

## Results

Compared to the long-term stability group, 135 DEGs were screened including 97 up-regulated and 38 down-regulated genes. We exhibited the top 7 significant terms of GOBP functional annotation enriched by Metascape (Fig □), and those terms were selected for the topic biological process (Table □) which had strong certainty (−log(p) >10) and direct relationship with skin development and differentiation. Most of these terms were identified as key processes of melanoma metastasis from a recent study^(11)^. Finally, 35 genes associated with the topic 7 GOBP terms were used to construct a PPI network (Fig □). The network involved 27 genes. ITGA6 was the only gene down-regulated in the rapid progression group, whereas the other genes were all up-regulated.

**Fig 1:**
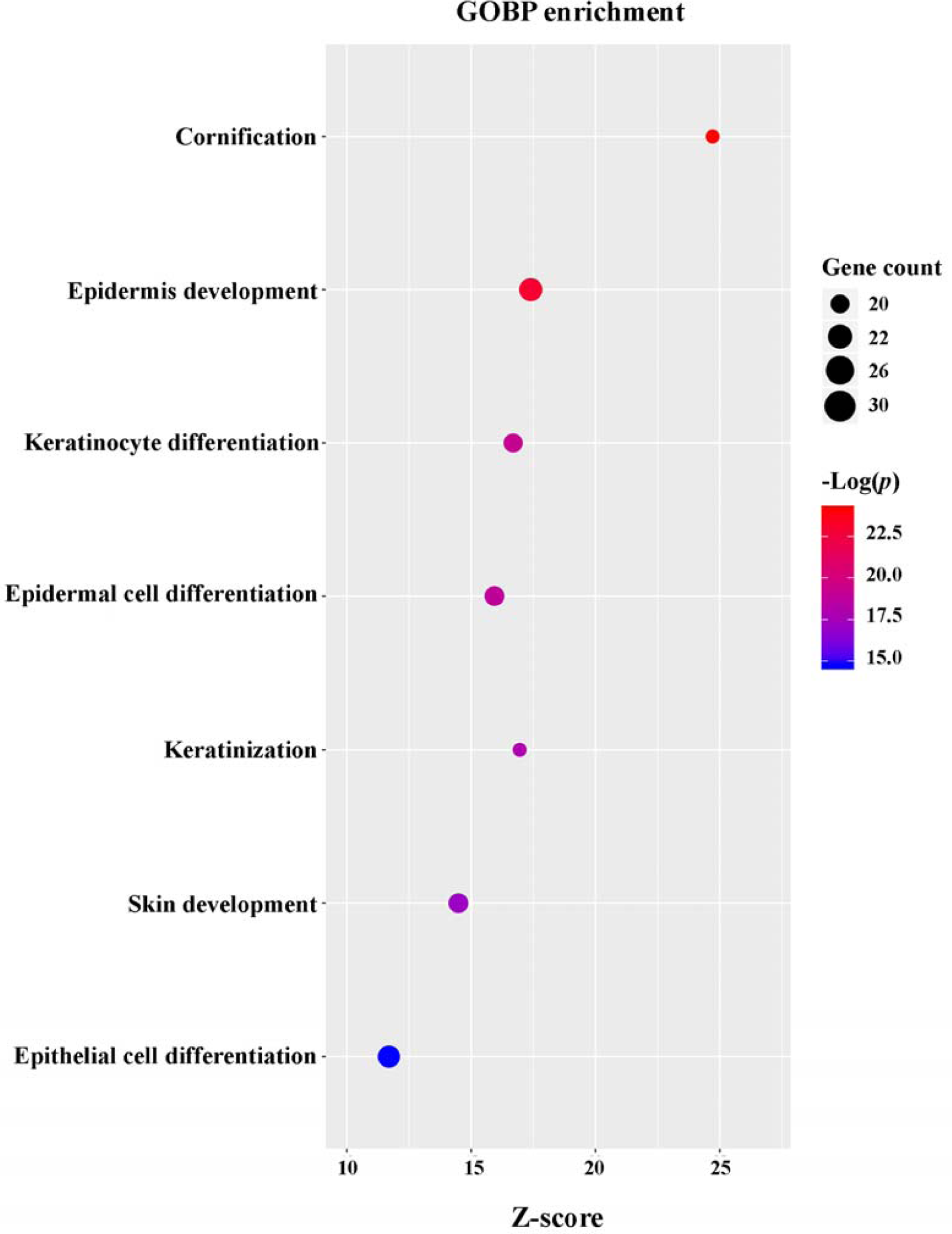
The top 7 significant terms of GOBP functional annotation enriched by Metascape

**Table 1:** Seven GOBP terms in DEGs.

| Name | GOBP Code | – Log(p) | Z-score | Gene Count |
| --- | --- | --- | --- | --- |
| Cornification | GO:0070268 | 24.13 | 24.71 | 20 |
| Epidermis Development | GO:0008544 | 22.94 | 17.40 | 30 |
| Keratinocyte Differentiation | GO:0030216 | 19.10 | 16.69 | 23 |
| Epidermal Cell Differentiation | GO:0009913 | 18.74 | 15.94 | 24 |
| Keratinization | GO:0031424 | 17.92 | 16.95 | 20 |
| Skin Development | GO:0043588 | 17.12 | 14.49 | 24 |
| Epithelial Cell Differentiation | GO:0030855 | 14.73 | 11.69 | 28 |

**Fig 2:**
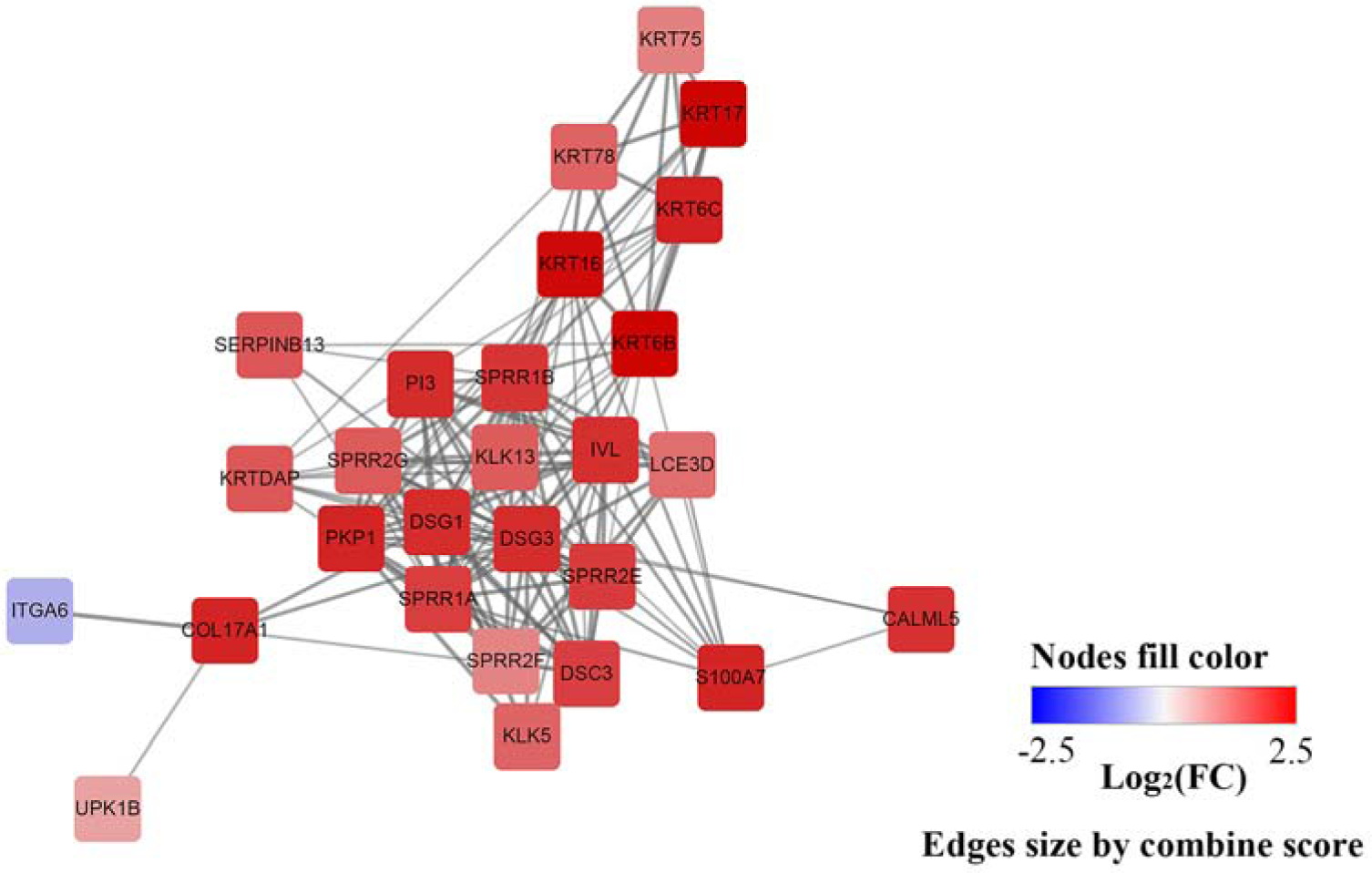
PPI network of the key functional gene (FC, fold-change of gene mRNA expression value). The combined score among the nodes was calculated by the STRING app

We randomly divided 372 patients for the prediction of disease progression into train and test groups (7:3 ratio). LASSO Cox regression model and 3-folds cross-validation were used to build a gene prognostic classifier in the train group (Fig □) which selected candidate 3 genes (COL17A1, ITGA6, and SPRR2F) from 27 key functional genes. According to the results of the lasso regression, we fitted a microarray data-related mathematical model for evaluating the disease progression as genetic risk score according to the following formula: 0.01547846 × log_2_ (COL17A1) − 0.09517359 × log_2_ (ITGA6) + 0.11394557 × log_2_ (SPRR2F). In order to evaluate the contribution of the genetic risk score in the Cox regression prediction model, we calculated the C-index value of the Cox regression model based on the genetic risk score, which increased the C-index value obviously as compared to the 27 key functional genes both in the train and test groups (Fig □). These results suggest that the genetic risk score is more effective and convenient in assessing disease prognosis than the inclusion of all the key functional genes.

**Fig 3:**
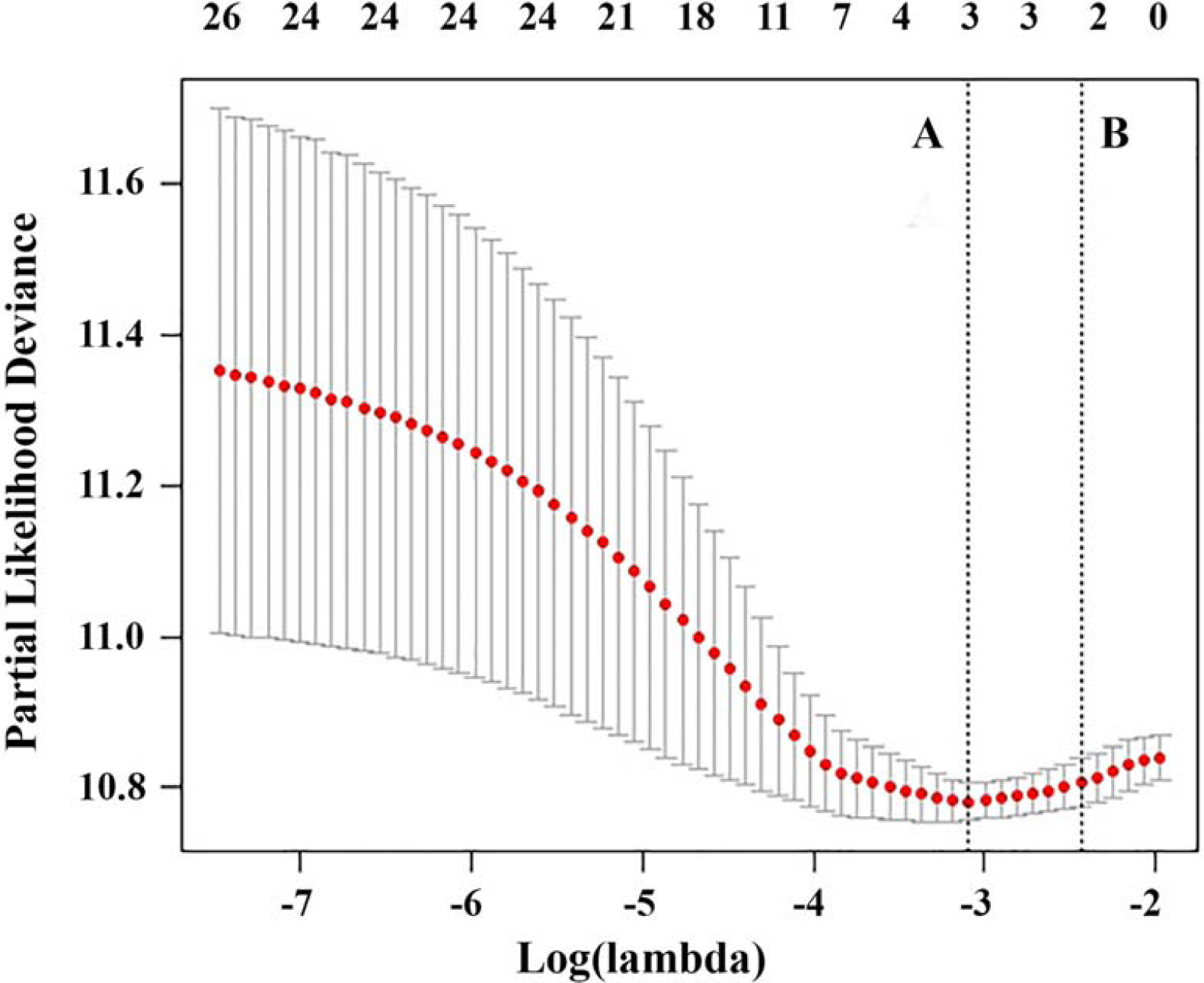
LASSO regression for selecting the contributive genes cross-validated by 3 folds. The A line refers to the min lambda value and the B line refers to the min lambda value plus 1 standard error

**Fig 4:**
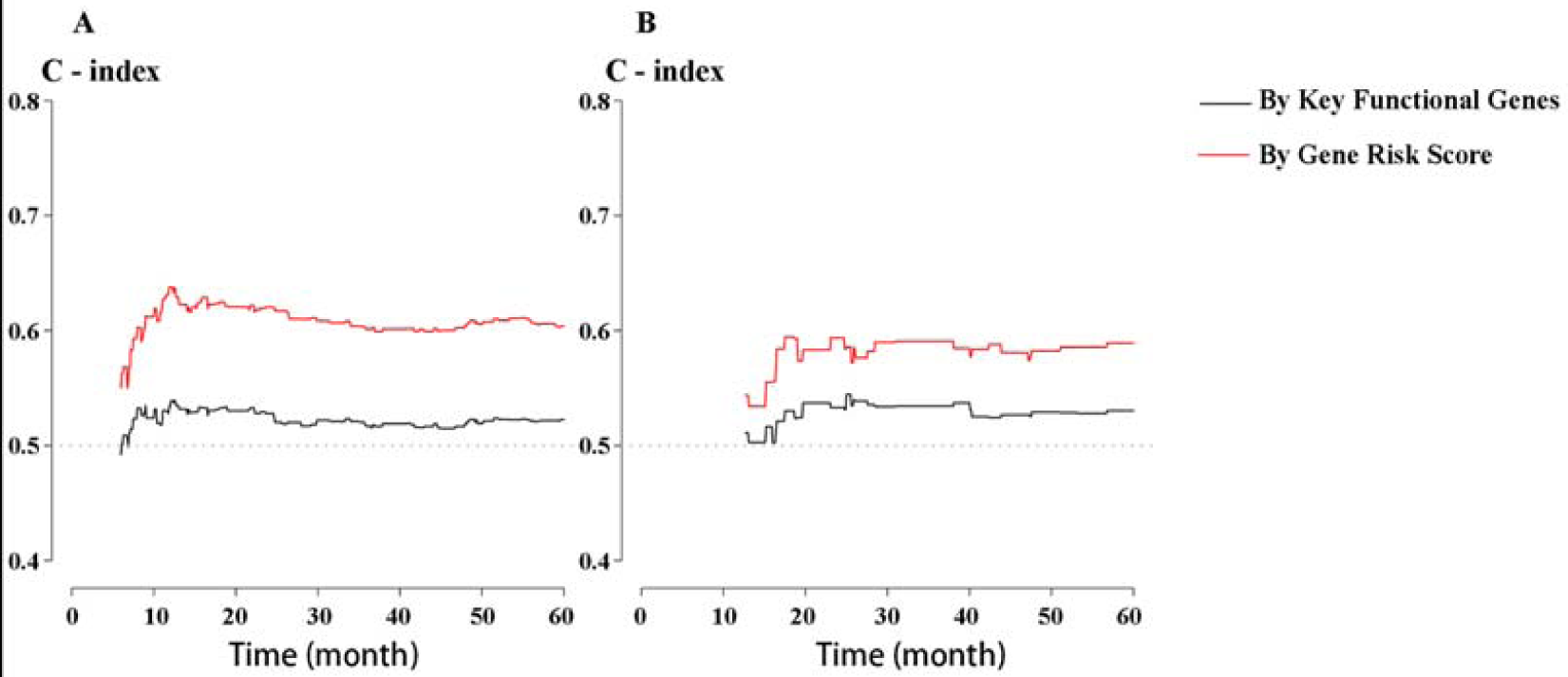
Sampling verification C-index of Cox regression model by bootstrap method 1000 step. A. train group, B. test group.

We combined the results of the train and test groups and then estimated the optimal cut-off value of the genetic risk score. We divided 372 patients into high-risk and low-risk groups by X-tile software (log-rank test p-value < 0.001). The cutoff value was **−** 0.73. Kaplan-Meier risk curves were constructed and the HR was calculated by univariate Cox regression analysis (Fig □). The high-risk score group was more prone to progression or cancer recurrence (HR, 2.39; 95% CI, 1.67**−**3.40; p < 0.001).

**Fig 5:**
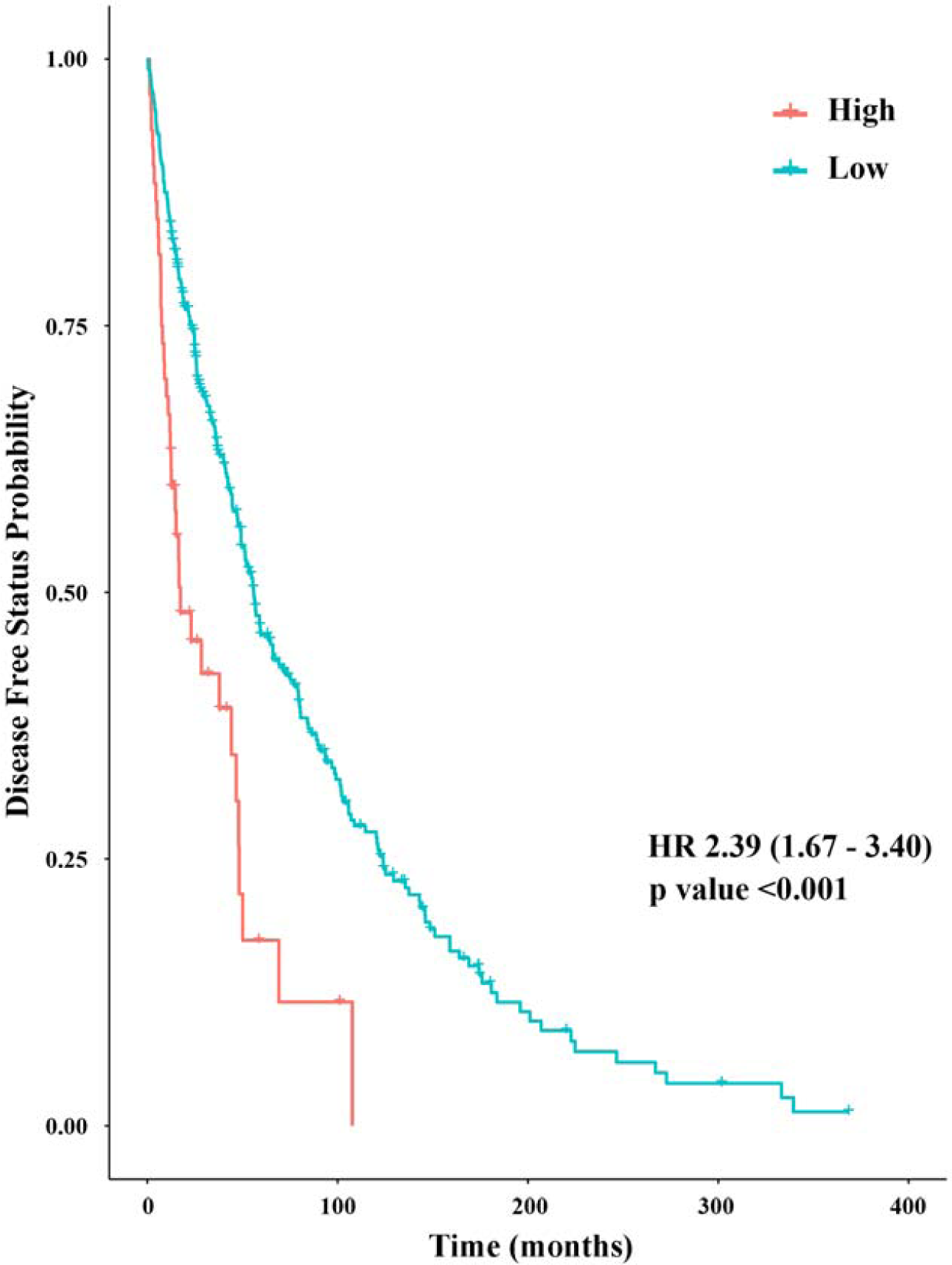
Kaplan-Meier curves of disease-free status survival according to the genetic risk score (HR, HR-hazard ratio calculated by univariate Cox regression).

Univariate Cox regression analysis was performed based on the clinical features in 372 patients. Age > 50 years, T stage > 2, lymph node metastasis, distant metastasis, and ulceration existence were found to be high-risk factors for disease progression. Age, lymph node metastasis, distant metastasis, and genetic risk score of high expression were found to be independent risk factors in the multivariate analysis combined with the genetic risk score (Table □). The C-index value of the multivariate model was 0.674. We calculated the IDI value for 1, 3, and 5 years disease-free probability prediction based on the clinical features model and genetic risk score associated model. It was found that the score of the combined model (IDI, 2.7%; p = 0.016) could weakly but significantly improve the estimation efficiency over the individual models (3.1%, and 3.1%, respectively; p = 0.034 and 0.014, respectively).

**Table 2:** Cox regression to predict disease progression.

| Subject | Univariate |  |  | Multivariate<sup>a</sup> |  |  |
| --- | --- | --- | --- | --- | --- | --- |
|  | HR | 95% CI | p | HR | 95% CI | p |
| Sex (male) | 1.19 | 0.91–1.54 | 0.198 | NA |  |  |
| Age (> 50 years) | 1.65 | 1.26–2.16 | < 0.001 | 1.55 | 1.07–2.24 | 0.022 |
| TNM-T (> 2) | 1.73 | 1.31–2.27 | < 0.001 | 1.09 | 0.75–1.57 | 0.652 |
| Lymph node (metastasis) | 2.21 | 1.69–2.89 | < 0.001 | 2.01 | 1.42–2.84 | < 0.001 |
| Distant metastasis (metastasis) | 2.20 | 1.32–3.66 | 0.002 | 2.25 | 1.08–4.67 | 0.030 |
| Ulceration (yes) | 1.84 | 1.35–2.50 | < 0.001 | 1.36 | 0.96–1.92 | 0.083 |
| Pharmaceutical (treated) | 0.85 | 0.62–1.16 | 0.292 | NA |  |  |
| Genetic risk score (high expression) | 2.39 | 1.67–3.40 | < 0.001 | 2.22 | 1.45–3.42 | < 0.001 |

*HR, hazard ratio; CI, confidence interval; NA, not available; a, complete information for age, T stage, N stage, M stage, ulceration, and genetic risk score was included in the multivariate Cox regression.*

The foregoing research has confirmed that the genetic risk score is an effective and independent marker in predicting the progression and recurrence of melanoma. Here, we explored the value of the genetic risk score in predicting the overall survival. A total of 425 patients (excluding 9 patients with incomplete survival information and 35 living patients with an observation time of < 1 year) were included in the TCGA melanoma cohort. Using the cut-off value of genetic risk score as previously predicted (**−** 0.73) as the boundary, these 425 patients were divided into two groups. Kaplan-Meier risk curves and univariate Cox regression analysis (Fig □) showed that the high-risk score group was more susceptible to death (HR, 2.92; 95% CI, 2.09-4.09; p < 0.001).

**Fig 6:**
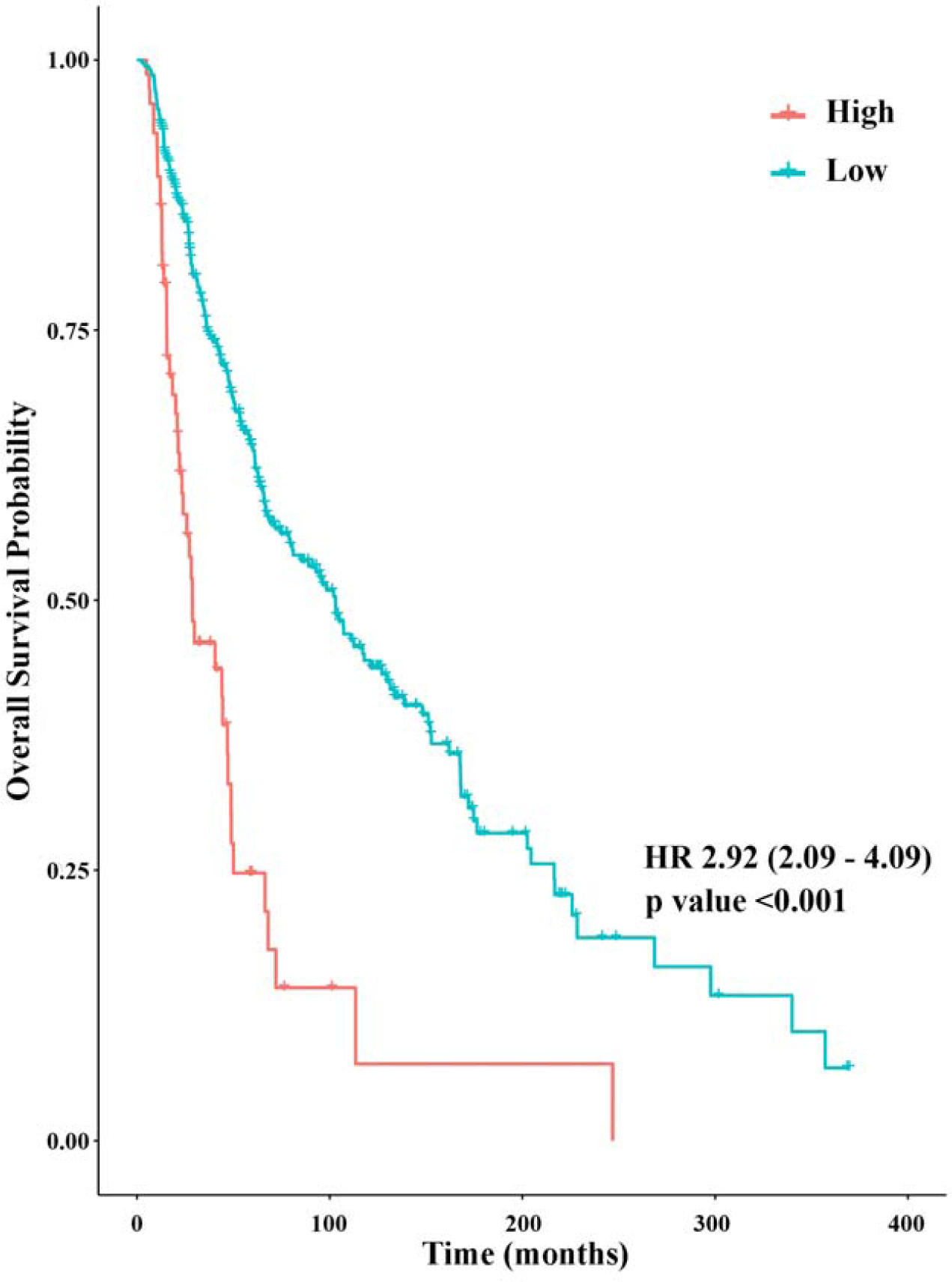
Kaplan-Meier curves of overall survival according to the genetic risk score. HR, hazard ratio calculated by univariate Cox regression.

Through univariate Cox regression analysis, we found that age of > 50 years, T stage of > 2, lymph node metastasis, and presence of ulceration were high-risk factors for death. By performing a multivariate analysis combined with the clinical risk features and genetic risk score, we identified that lymph node metastasis, presence of ulceration, and high expression in the genetic risk score are independent and effective risk factors for estimating the overall survival rate (Table □). The C-index value of the multivariate model was 0.684. The IDI value in 1, 3, and 5 years were calculated among the independent clinical features with or without genetic risk score. The results suggest that in one year, the genetic risk score is rarely effective in improving the model prediction ability (IDI, 0.8%; p = 0.555), whereas, in 3 or 5 years, the improvement in the genetic risk score could significantly improve the estimation efficiency (IDI 5.3% and 5.8%, respectively; p = 0.008 and < 0.001, respectively).

**Table 3:** Cox regression to predict the overall survival.

| Subject | Univariate |  |  | Multivariate<sup>a</sup> |  |  |
| --- | --- | --- | --- | --- | --- | --- |
| | HR | 95% CI | $p$ | HR | 95% CI | $p$ |
| Sex (male) | 1.14 | 0.86–1.52 | 0.349 | NA |  |  |
| Age (> 50 years) | 1.67 | 1.25–2.23 | <0.001 | 1.42 | 0.96–2.12 | 0.083 |
| TNM-T (> 2) | 1.98 | 1.46–2.68 | <0.001 | 1.24 | 0.83–1.84 | 0.294 |
| Lymph node (metastasis) | 1.75 | 1.31–2.34 | <0.001 | 1.82 | 1.23–2.65 | 0.002 |
| Distant metastasis (metastasis) | 1.74 | 0.92–3.29 | 0.089 | NA |  |  |
| Ulceration (yes) | 2.11 | 1.51–2.94 | <0.001 | 1.59 | 1.10–2.32 | 0.015 |
| Pharmaceutical (treated) | 0.86 | 0.62–1.20 | 0.369 | NA |  |  |
| Genetic risk score (high expression) | 2.92 | 2.09–4.09 | <0.001 | 2.72 | 1.76–4.21 | <0.001 |

HR, hazard ratio; CI, confidence interval; NA, not Available; a, complete information for age, T stage, N stage, M stage, ulceration and genetic risk score were included in the multivariate Cox regression.

## Discussion

Previous studies have identified several genetic markers associated with the outcome of melanoma^(12)^, such as BRAF, GNA11, BAP1, EIF1AX, SF3B1, and NFkB, most of which were associated with ubiquitous signaling pathways in the cells, such as MAPK, PI3K/AKT, and telomerase. Several melanoma gene-targeted drugs have been listed to be mostly involved in the immune-checkpoint pathways^(13-15)^. The seven topic biological processes selected were concerned with the development, differentiation, and programming of derma cell, which might have better functional specificity in the study of skin tumors. A total of 27 functional genes related to the 7 biological processes were found by the PPI network analysis. We believed that those genes would play important roles in melanoma progression and might become new targets of melanoma gene therapy.

To simplify the clinical application process, we used regression method to select 3 marker genes that had significant contribution in predicting the outcome of melanoma and developed a novel prognostic method based on the expression of these marker genes. Although no study has confirmed that the expression of these genes was directly associated with the outcomes of melanoma, some studies could corroborate our conjecture. COL17A1 codes collagen type XVII alpha 1 chain. Krenacs’s^(16)^ has suggested that the accumulation of COL17 endodomain in the melanocytes was associated with malignant transformation of melanoma. In another bioinformatics research^(17)^ based on an independent data source, the upstream lncRNA (RP11-361L15.4) of COL17A1 was screened as a biomarker of melanoma. The results confirmed that COL17A1 is a key gene in the development and progression of melanoma. ITGA6 is coded by the integrin subunit alpha 6. Lenci’s^(18)^ has analyzed multiple members of the ITGA gene family, and the variants of ITGA6 were associated with melanoma susceptibility. Karhemo^(19)^ considered that ITGA6 plays an important role in the metastasis of melanoma. Further study^(20)^ have provided evidence that the genes enriched in ITGA6-B4 pathway were downregulated in the invasive primary melanoma-derived clones. SPRR2F codes small proline-rich protein 2F. SPRR2F was identified as a targeted gene^(21)^ in epithelial malignant tumors and some other members of the SPRR protein family, which were considered to augment epithelial proliferation and initiation of malignancy^(22)^. Furthermore, SPRRs were also up-regulated after UV exposure^(23)^, which indicated that this gene family might mediate some pathological changes in causing skin phototoxicity and may also play a crucial role in melanoma progression. Based on these studies along with the results of our study, it is reasonable to use genetic risk score to evaluate the outcome of melanoma.

For the prediction of disease progression or overall survival, the genetic risk score was found to be an independent and effective risk factor. The results of the IDI calculation showed that as compared to the Cox regression model developed based on clinical features, the genetic risk score combined model could improve the effectiveness of the prediction. We suggested that our novel method has promising potential for clinical application not only in disease prediction but also in gene-targeted therapy. But unfortunately, the C-index value of the Cox regression model seemed to be little weak, and we did not perform an appropriate external validation. These two drawbacks might limit our findings for clinical application and warrant further research.

## Acknowledgements

The authors would like to thank the BBS.DXY.CN for the inspiration of this research.

## Availability of data and materials

All data generated or analyzed during this study are included in this published article.

## Author’s contributions

HGH was the major contributor in designing the research, performing bioinformatic and statistical analysis and writing the manuscript. GQ collated and analyzed the clinical data form TCGA database. LXH supervised the study. All authors have read and approved the final manuscript.

## Ethics approval and consent to participate

Not applicable.

## Consent for publication

Not applicable.

## Reference

1. Ferlay J, Soerjomataram I, Dikshit R, Eser S, Mathers C, Rebelo M, Parkin DM, Forman D, Bray F: Cancer incidence and mortality worldwide: sources, methods and major patterns in GLOBOCAN 2012. Int J Cancer 2015, 136:E359–386.

2. Siegel RL, Miller KD, Jemal A: Cancer statistics, 2019. CA Cancer J Clin 2019, 69:7–34.

3. Gershenwald JE, Scolyer RA, Hess KR, Sondak VK, Long GV, Ross MI, Lazar AJ, Faries MB, Kirkwood JM, McArthur GA, et al: Melanoma staging: Evidence-based changes in the American Joint Committee on Cancer eighth edition cancer staging manual. CA Cancer J Clin 2017, 67:472–492.

4. Shain AH, Bastian BC: From melanocytes to melanomas. Nat Rev Cancer 2016, 16:345–358.

5. Chandran UR, Medvedeva OP, Barmada MM, Blood PD, Chakka A, Luthra S, Ferreira A, Wong KF, Lee AV, Zhang Z, et al: TCGA Expedition: A Data Acquisition and Management System for TCGA Data. PLoS One 2016, 11:e0165395.

6. Cerami E, Gao J, Dogrusoz U, Gross BE, Sumer SO, Aksoy BA, Jacobsen A, Byrne CJ, Heuer ML, Larsson E, et al: The cBio cancer genomics portal: an open platform for exploring multidimensional cancer genomics data. Cancer Discov 2012, 2:401–404.

7. Kohl M, Wiese S, Warscheid B: Cytoscape: software for visualization and analysis of biological networks. Methods Mol Biol 2011, 696:291–303.

8. Szklarczyk D, Franceschini A, Wyder S, Forslund K, Heller D, Huerta-Cepas J, Simonovic M, Roth A, Santos A, Tsafou KP, et al: STRING v10: protein-protein interaction networks, integrated over the tree of life. Nucleic Acids Res 2015, 43:D447–452.

9. Tibshirani R: The lasso method for variable selection in the Cox model. Stat Med 1997, 16:385–395.

10. Camp RL, Dolled-Filhart M, Rimm DL: X-tile: a new bio-informatics tool for biomarker assessment and outcome-based cut-point optimization. Clin Cancer Res 2004, 10:7252–7259.

11. Wang B, Qu XL, Chen Y: Identification of the potential prognostic genes of human melanoma. J Cell Physiol 2019, 234:9810–9815.

12. Rabbie R, Ferguson P, Molina-Aguilar C, Adams DJ, Robles-Espinoza CD: Melanoma subtypes: genomic profiles, prognostic molecular markers and therapeutic possibilities. J Pathol 2019, 247:539–551.

13. Leonardi GC, Falzone L, Salemi R, Zanghi A, Spandidos DA, McCubrey JA, Candido S, Libra M: Cutaneous melanoma: From pathogenesis to therapy (Review). Int J Oncol 2018, 52:1071–1080.

14. Bhandaru M, Rotte A: Monoclonal Antibodies for the Treatment of Melanoma: Present and Future Strategies. Methods Mol Biol 2019, 1904:83–108.

15. Mason R, Au L, Ingles Garces A, Larkin J: Current and emerging systemic therapies for cutaneous metastatic melanoma. Expert Opin Pharmacother 2019, 20:1135–1152.

16. Krenacs T, Kiszner G, Stelkovics E, Balla P, Teleki I, Nemeth I, Varga E, Korom I, Barbai T, Plotar V, et al: Collagen XVII is expressed in malignant but not in benign melanocytic tumors and it can mediate antibody induced melanoma apoptosis. Histochem Cell Biol 2012, 138:653–667.

17. Zhang Q, Wang Y, Liang J, Tian Y, Zhang Y, Tao K: Bioinformatics analysis to identify the critical genes, microRNAs and long noncoding RNAs in melanoma. Medicine (Baltimore) 2017, 96:e7497.

18. Lenci RE, Rachakonda PS, Kubarenko AV, Weber AN, Brandt A, Gast A, Sucker A, Hemminki K, Schadendorf D, Kumar R: Integrin genes and susceptibility to human melanoma. Mutagenesis 2012, 27:367–373.

19. Karhemo PR, Ravela S, Laakso M, Ritamo I, Tatti O, Makinen S, Goodison S, Stenman UH, Holtta E, Hautaniemi S, et al: An optimized isolation of biotinylated cell surface proteins reveals novel players in cancer metastasis. J Proteomics 2012, 77:87–100.

20. Vizkeleti L, Kiss T, Koroknai V, Ecsedi S, Papp O, Szasz I, Adany R, Balazs M: Altered integrin expression patterns shown by microarray in human cutaneous melanoma. Melanoma Res 2017, 27:180–188.

21. Contreras CM, Akbay EA, Gallardo TD, Haynie JM, Sharma S, Tagao O, Bardeesy N, Takahashi M, Settleman J, Wong KK, Castrillon DH: Lkb1 inactivation is sufficient to drive endometrial cancers that are aggressive yet highly responsive to mTOR inhibitor monotherapy. Dis Model Mech 2010, 3:181–193.

22. Carregaro F, Stefanini AC, Henrique T, Tajara EH: Study of small proline-rich proteins (SPRRs) in health and disease: a review of the literature. Arch Dermatol Res 2013, 305:857–866.

23. Cho BA, Yoo SK, Seo JS: Signatures of photo-aging and intrinsic aging in skin were revealed by transcriptome network analysis. Aging (Albany NY) 2018, 10:1609–1626.

